# Early life stress impairs VTA coordination of BLA network and behavioral states

**DOI:** 10.1101/2023.09.16.558081

**Authors:** Bradly T. Stone, Pantelis Antonoudiou, Eric Teboul, Garrett Scarpa, Grant Weiss, Jamie L. Maguire

## Abstract

Motivated behaviors, such as social interactions, are governed by the interplay between mesocorticolimbic structures, such as the ventral tegmental area (VTA), basolateral amygdala (BLA), and medial prefrontal cortex (mPFC). Adverse childhood experiences and early life stress (ELS) can impact these networks and behaviors, which is associated with increased risk for psychiatric illnesses. While it is known that the VTA projects to both the BLA and mPFC, the influence of these inputs on local network activity which govern behavioral states – and whether ELS impacts VTA-mediated network communication – remains unknown. Our study demonstrates that VTA inputs influence BLA oscillations and mPFC activity, and that ELS weakens the ability of the VTA to coordinate BLA network states, likely due to ELS-induced impairments in dopamine signaling between the VTA and BLA. Consequently, ELS mice exhibit increased social avoidance, which can be recapitulated in control mice by inhibiting VTA-BLA communication. These data suggest that ELS impacts social reward via the VTA-BLA dopamine network.

## Introduction

Adverse childhood experiences (ACEs) are major risk factors for psychiatric illnesses. For example, maternal neglect is known to impair cognition, weaken resiliency towards future stressors, and increase the propensity to develop mood and substance use disorders in affected offspring ([1, 2], for review see [3]). Recent reports show that roughly 20% of children in the United States are exposed to ACEs [4] and research has found a striking comorbidity between frequency and severity of ACEs with psychiatric illnesses [5–7]. In conjunction with these findings, decades of work have implicated the mesocorticolimbic system (MCL) as being a psychophysiological focal point due to its susceptibility to the effects of stressors [8, 9], including early life stress (ELS; [10, 11]), and its involvement in proper establishment and maintenance of reward processing and subsequent motived behaviors [12]. These epidemiological and deductive research studies support a role for the MCL in contributing to behavioral deficits associated with early life adversity; however, the mechanisms through which ELS influences the MCL to induce altered behavioral outcomes remains unclear. Revealing these mechanisms has the potential to provide both useful information regarding the pathophysiology of disease as well as potential novel avenues for therapeutic intervention.

Many aspects of motivation involve dopaminergic (DA) neurons within and across the MCL. Further, studies have shown that the MCL DA system is not fully developed until adulthood in both humans [13] and rodents [14], making adolescents particularly vulnerable to ACEs and ELS, respectively [15]. Specifically, in rodents, the MCL DA system hits maturity on postnatal day 21 (PND21, [14]) with hindbrain features (e.g. neurons and neuropeptides) known to be structurally and functionally altered following ELS by this timepoint [2, 16]. While a growing body of research has narrowed the impact of ELS to a subset of nodes within the MCL, the mechanisms mediating the developmental impact of altered DA signaling on emotional processing and motivation are not fully understood.

Accumulating evidence from our lab and others demonstrates that network states (distinct oscillatory activity, quantified as local field potentials; LFPs) in the basolateral amygdala (BLA) govern behavioral states [17–19], particularly those involved in valence processing [20, 21]. Moreover, the BLA is well established as the primary emotional hub within the mammalian brain [22–24]. Previous research has also shown that the VTA [25, 26] and connections between the VTA and regions across the MCL [27–29] are necessary for proper regulation of incentive attributions of salience to stimuli, which influence behaviors. The VTA has direct projections to the BLA [30, 31], which monosynaptically projects to the medial prefrontal cortex [32–34] (mPFC; anterior region of the MCL) – a region also known to be susceptible to stress [35, 36] and is involved in motivation processing [37–39]. Therefore, it is reasonable to hypothesize that VTA inputs into the BLA likely influence motivated behaviors.

Despite the acknowledgment of the VTA’s projections to both the BLA and mPFC, the specific influence of these DA pathways on local network dynamics and motivated behaviors remains elusive. Moreover, the impact of ELS on VTA-BLA communication and the mechanisms mediating the subsequent behavioral consequences have yet to be fully elucidated. This study addresses these gaps in knowledge by examining the intricate modulatory role of the VTA in regulating oscillatory states in the BLA and, in turn, influencing mPFC activity and social motivation. We show that the VTA robustly controls BLA and entrains mPFC network states, driving BLA-mPFC coherence, while ELS diminishes these effects. Further, our results suggest that this dampening of VTA-mediated information processing in the BLA and routing through the mPFC following ELS involves a reduction in both the quantity of BLA-directed DA projections from VTA, as well as BLA DA sensitivity. We go on to show that activation of VTA projections to the BLA following ELS results in social avoidance, a phenotype only recapitulated in control mice following inhibition of these VTA efferents, further supporting this network’s involvement in stress vulnerability, and its control over stimulus incentive assignment. These findings have implications for our understanding of the etiology and potential therapeutic interventions for psychiatric disorders marked by disrupted motivation and reward processing.

## Results

To determine whether VTA to BLA (VTA_BLA_) activation influences BLA network activity, and if developmental stress alters these responses, we implemented an early life stress (ELS) protocol in C57BL/6J mice, such that mice were either standard-reared (CNT) or subjected to maternal separation stress for 3 hours daily, 5 days per week from PND1 – PND21 (**Figure 1A**). Following weaning and maturation, infusion of AAV-ChR2-tdT viral vector into VTA resulted in expression of ChR2-tdT in projection neurons from VTA, with strong terminal expression centered in BLA (**Figure 1B**). Pulsed red light (640 nm) photoexcitation (5-40 Hz) in the BLA produced robust phase-locked LFP responses in both the BLA and mPFC (**Figure 1C; Supp1A-D**), indicating that excitation of VTA inputs into the BLA is sufficient to control BLA network synchronization across a range of frequencies and entrain the mPFC network, with ELS attenuating this effect (**Figure 1D, Stats Table 1**). These results suggest that the ability for VTA inputs into the BLA to coordinate BLA-mPFC network states is impaired following ELS, which may give rise to aberrant processing of salient cues [40, 41] and, thus, maladaptive behaviors ([42], for review see [43]).

**Figure 1:**
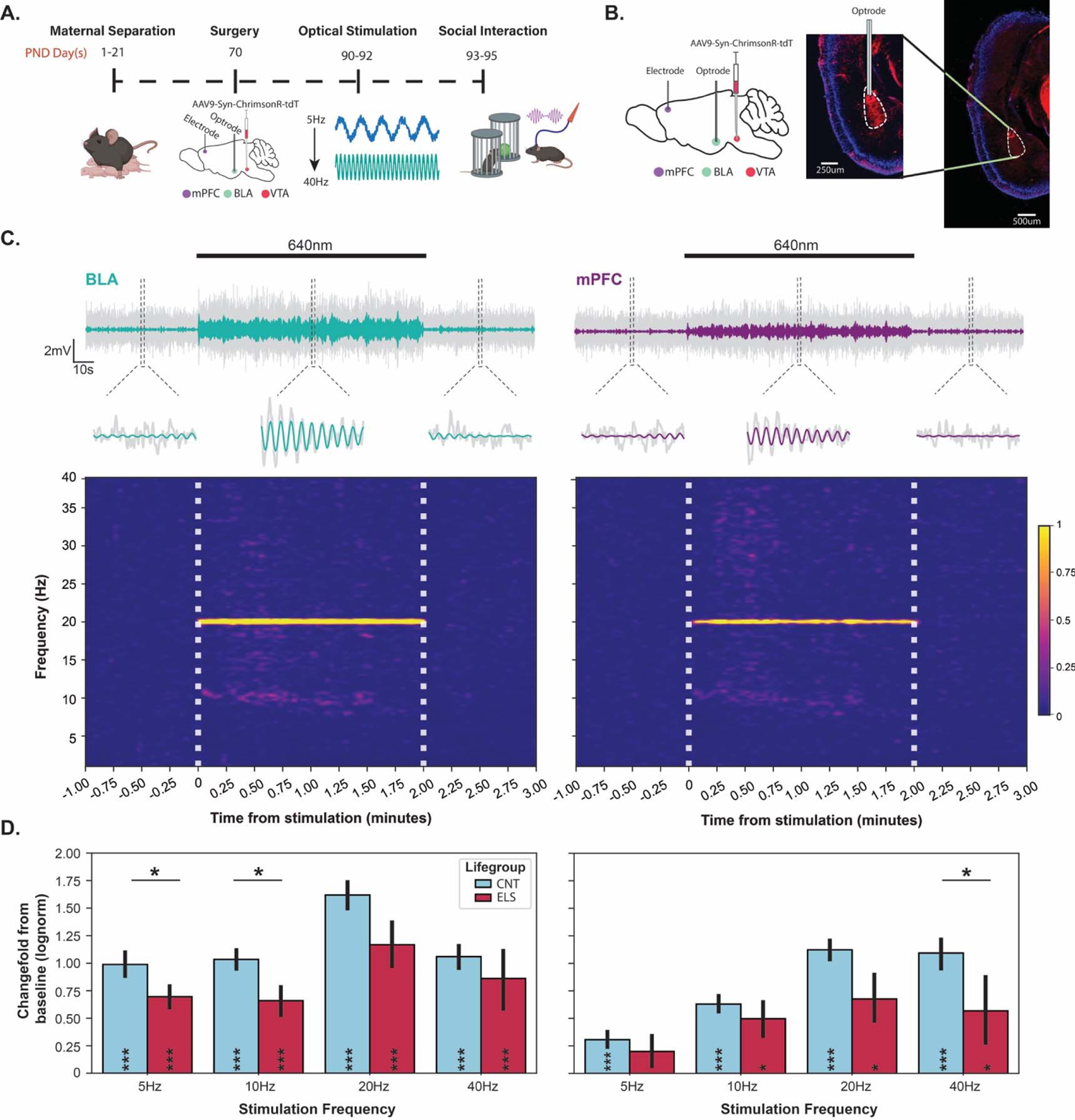
ELS inhibits ability for VTA_BLA_ to drive network activity. (**A**) Experimental timeline for ELS cohort. Timeline is identical for CNT cohort excluding maternal separation (D1-21). (**B**) Left: Schematic of sagittal view shows at PND70, AAV-Syn-ChrimsonR-tDT was injected into VTA, with an optrode and electrode placed into BLA and mPFC respectively; Right: Representative fluorescence images of ChR2-tdT expression in the BLA. (**C**) Top: Representative 20Hz LFP trace (colored) overlaid on raw LFP trace (2-100Hz; grey) from BLA (left) and mPFC (right); Middle: Zoomed in portions from above traces show 5 periods before, during, and after 10mW 640nm photoexcitation; Bottom: Wavelet transformation from respective traces; vertical dashed white lines indicate onset/offset of 640nm laser. (**D**) Summary of LFP power ratio from BLA (left) and mPFC (right) between pulsed stimulation and baseline periods (baseline = 0) where power was measured at each stimulation frequency for CNT (blue, n = 7) and ELS (red, n = 8) conditions. Asterisks within bars indicate significant change from baseline. Asterisks atop horizontal bars indicate difference between CNT and ELS. * *p* < 0.05, ** *p* < 0.01, *** *p* < 0.001. Error bars represent SEM.

To examine the impact of ELS on VTA_BLA_-driven BLA-mPFC coordinated activity, we computed the phase-locking value (PLV) across a series of photoexcitation frequencies (**Figures 2A-D**). As expected, photoactivation of VTA_BLA_ terminals promotes BLA activity to lead the activity in the mPFC (**Figures 2B** and **2D, Stats Table 2**). Further, our results reveal an attenuation in the VTA-mediated coherence between BLA and mPFC following ELS (**Figure 2E).** Specifically, VTA_BLA_ activation significantly enhances BLA-mPFC coherence in all frequencies greater than 5Hz, regardless of developmental stress. However, 40Hz stimulation-induced phase coherence in the ELS group is significantly reduced relative to controls (**Figure 2E, Stats Table 2**)—a frequency previously shown to modulate sociability (i.e. interaction and avoidance behaviors) in rodents [44, 45].

**Figure 2:**
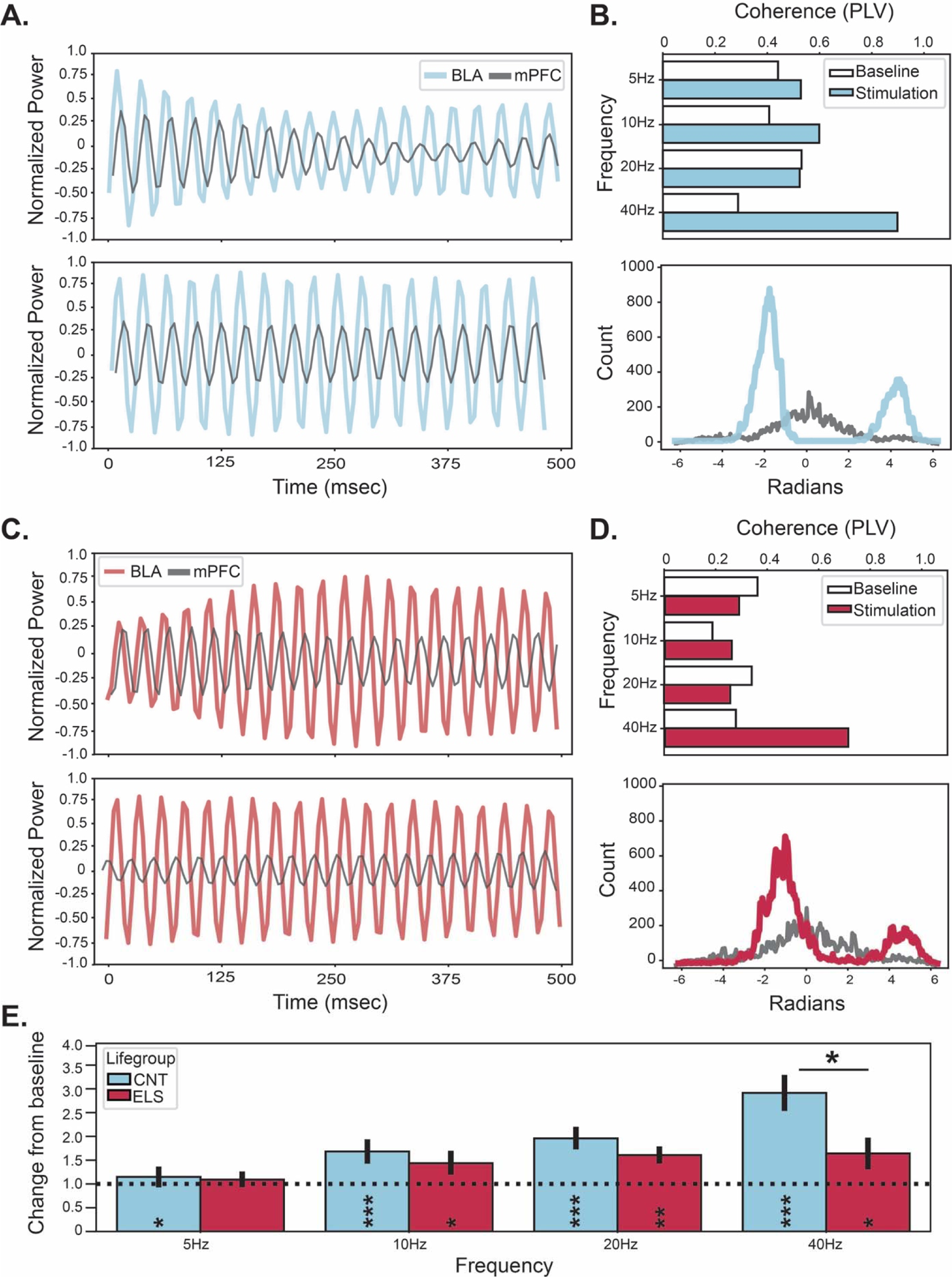
ELS minimizes VTA_BLA_ →mPFC functional network connectivity. (**A**) Representative LFP traces from BLA and mPFC before (Top) and during (Bottom) 40Hz VTA_BLA_ photoexcitation for a CNT mouse. (**B**) Top: Representative summary bar plots for animal in (**A**) detailing VTA evoked BLA-mPFC coherence at each stimulation frequency; Bottom: Phase histogram for baseline (grey) and 40Hz stimulation (red) in (**A**). (**C-D**) Same analyses seen in (**A-B**) for a representative ELS mouse. (**E**) Summary of LFP coherence ratio between pulsed stimulation and baseline periods where BLA-mPFC coherence is measured at each stimulation frequency for CNT (blue, n = 7) and ELS (red, n = 8) conditions. Dashed horizontal line indicates baseline response. Asterisks within bars indicate significant change from baseline. Asterisks atop horizontal bars indicate difference between CNT and ELS. * *p* < 0.05, ** *p* < 0.01, *** *p* < 0.001. Error bars represent SEM.

Given our observation that ELS diminishes the influence of VTA inputs into the BLA, stifling the coherent signaling between the BLA and mPFC, we hypothesized that these phenomena arise from 1) cellular/synaptic changes; 2) neuroplastic/structural changes; 3) impaired neurotransmitter signaling/sensitivity; or 4) a combination of the three during this developmental critical period [15, 46].

To interrogate whether ELS induces functional deficits in synaptic signaling between the VTA and BLA, we performed whole-cell patch-clamp recordings on BLA principal neurons to measure DA sensitivity. We first determined the basic intrinsic membrane and action potential (AP) properties (**Supp 2**) following an input-output (IO) protocol before (baseline) and after bath application of a highly potent and selective D1-receptor [D1R] antagonist (10-nM SCH 23390; SCH) (**Figure 3A**). We observed no differences in baseline waveform shape, AP frequency, nor resting membrane potential (RMP) following ELS (**Figure 3B** & **Supp2**). However, a D1R-block caused depolarization of BLA principal cells (**Figures 3C-D, Stats Table 3**) and also suppressed high-input firing (observed at 80% vs 20% input) in CNT, but not ELS animals (**Figure 3E, Stats Table 3**).

**Figure 3:**
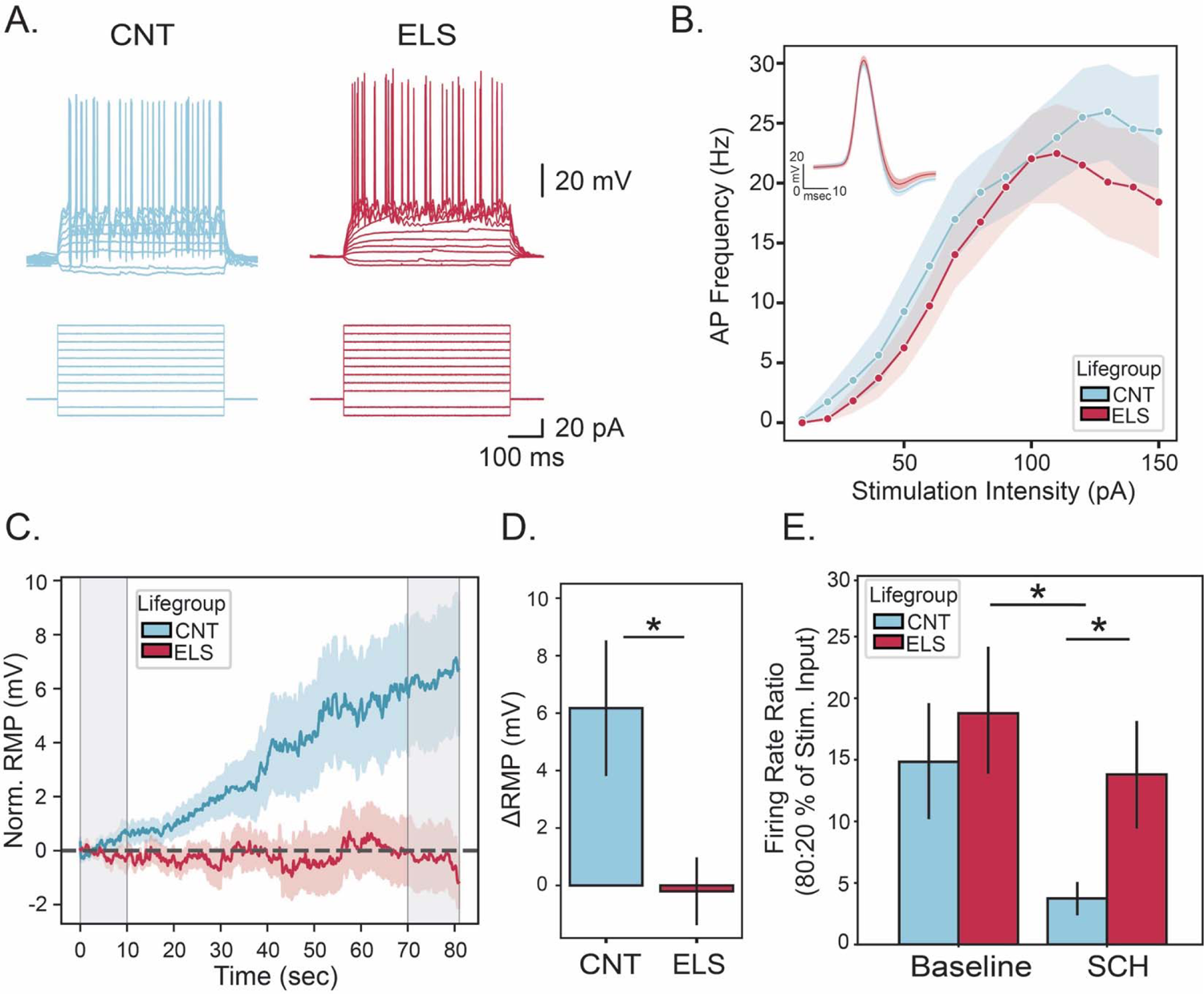
ELS impairs D1-mediated activity in BLA principal cells. (**A**) Representative traces show firing rate responses to rectangular current injections for CNT and ELS principal BLA neurons. (**B**) Inlay: Mean waveforms from CNT (N = 5 animals, n = 13 cells) and ELS (N = 5 animals, n = 12 cells) BLA neurons; Main: Summary plot of action potential (AP) frequency measured at each stimulation intensity for CNT and ELS BLA neurons. (**C**) Summary plot of normalized resting membrane potential (Norm. RMP) for CNT (N = 5 animals, n = 9 cells) and ELS (N = 5 animals, n = 7 cells) treatment-paired cells from (B) following bath application of 10-nM SCH 23390 (SCH). (**D**) Summary plot showing difference in Norm. RMP between the first and last 10secs (shown in grey in [**C**]) of SCH application. (**E**) Grouped bar plot shows firing rate ratio (FR Ratio) between AP firing observed at 80% and 20% input before and after D1R-block. * *p* < 0.05. Error bars and shading represent SEM.

To investigate the potential role of developmental stress on structural plasticity leading to impaired DA signaling in the BLA (**Figure 3**), we questioned whether ELS alters the machinery necessary for DA signaling. We injected CNT and ELS animals with rAAV-GCamP6f-tdT into BLA (**Figure 4A**), a viral approach that retrogradely labels VTA_BLA_ projecting neurons. Following adequate time for viral expression, we performed immunohistochemistry to quantify changes wrought by ELS in both the BLA and VTA. Given that D1Rs are 1) the most abundant DA receptor within BLA [47, 48], 2) known to be susceptible to stressors [49, 50], and 3) our findings of different effects of a D1R antagonist on BLA principal neuron activity (**Figure 3**), our initial investigation tested the abundance of BLA D1Rs. We found that ELS did not change the quantity of DAPI-expressing cells (**Figure 4B, Stats Table 4**), nor the density of D1Rs in BLA (**Figure 4C, Stats Table 4**), suggesting that the developmental impact of ELS likely originates in the hindbrain. To specifically investigate the influence of ELS on hindbrain structures, we hypothesized that ELS-induced differences should be constrained to the VTA, rather than the substantia nigra (SN) – a neighboring region known to have less DA cells and to experience resiliency towards developmental stressors [51, 52]. The number of retrogradely-labeled VTA neurons projecting to the BLA (tdT+ cells) was not impacted by ELS for the SN, nor the VTA (**Figures 4D-E**). While the total number of TH+ cells was significantly higher in the VTA than the SN, increasing distal from bregma, ELS had no impact on these quantities (**Figures 4D-E and Supp2**). However, ELS did significantly reduce the number of TH+ neurons projecting from the VTA to the BLA (tdT+/TH+ cell colocalization) with no change in the number of TH+ neurons projecting from the SN to the BLA (**Figures 4D** and **F, Stats Table 4**). Therefore, ELS reduces the number of VTA_BLA_ DA projecting cells, suggesting that network architecture supporting DA signaling is impaired after early life stress.

**Figure 4:**
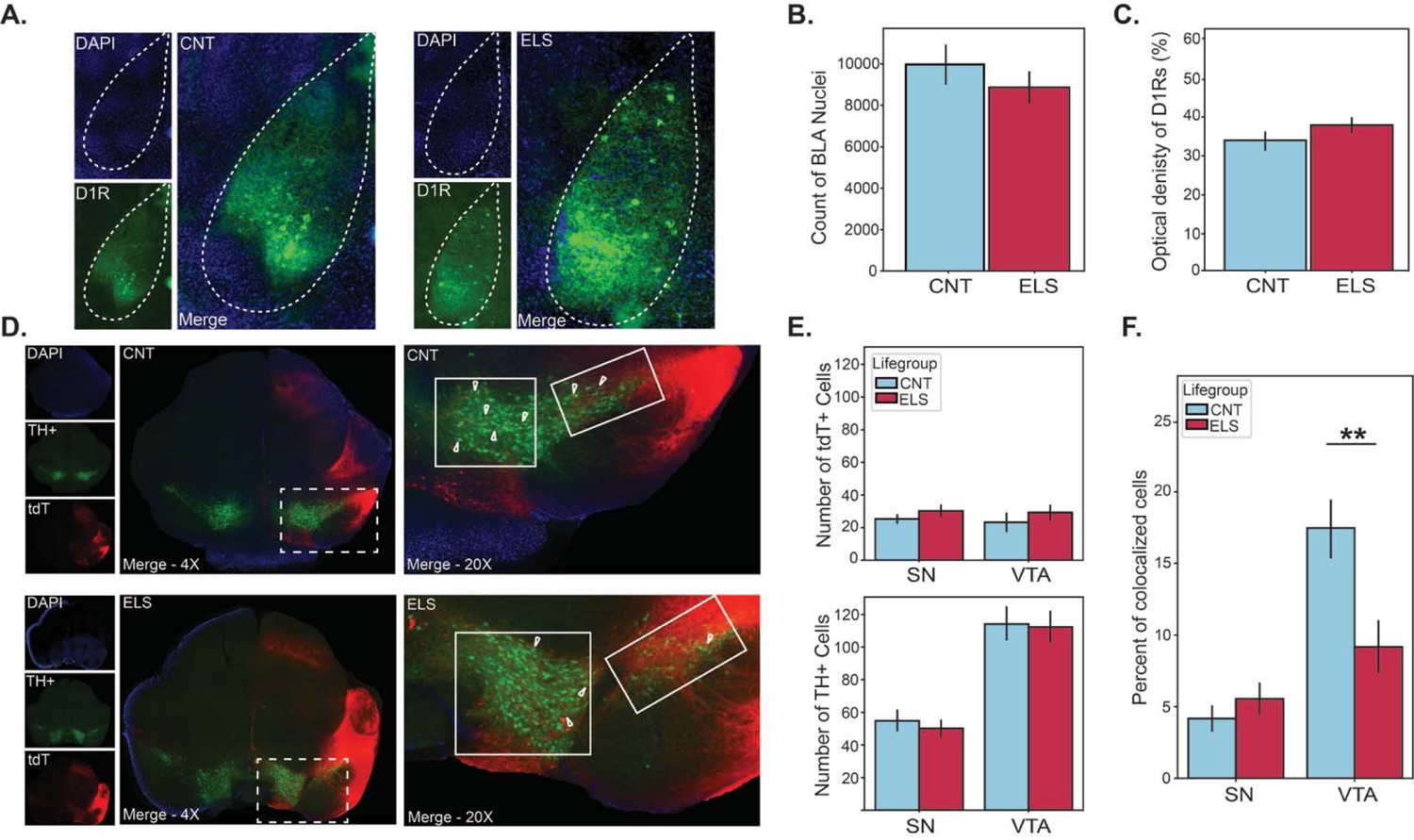
ELS reduces VTA→BLA dopaminergic inputs without impacting BLA D1 receptor quantity. (**A**) Representative 20X image for CNT (Left) and ELS (Right) BLA slice, outlined in white. DAPI: blue, D1 receptor (D1R) immunoreactivity: green. (**B**) Summary plot shows mean BLA nuclei count in CNT (N = 5 animals, n = 6 slice per animal) and ELS (N = 4 animals, n = 6 slice per animal). (**C**) Summary plot shows mean optical density of BLA D1Rs for CNT and ELS animals. (**D**) Left: Representative images of coronal hindbrain slices for a CNT (Top) and ELS (Bottom) animal at 4X. Right: 20x images of outlined rectangles on 4X images with arrow-tips indicating tdT-TH+ colocalized neurons; VTA: white square; substantia nigra (SN): white rectangle. (**E**) Top: Grouped bar plot shows average number of BLA-projecting neurons (tdT+) from SN and VTA. Bottom: Grouped bar plot shows average number of TH+ neurons in SN and VTA. (**F**) Grouped bar plot shows percentage of tdT-TH+ colocalized neurons in SN and VTA. ** *p* < 0.01. Error bars represent SEM.

Collectively, these results suggest that ELS causes a reduction in VTA_BLA_ DA release. To test this, we measured BLA DA repsonses by virally expressing a DA fluorescent sensor, GRABDA_2M_ [53], in the BLA of animals outfitted for microendoscopic imaging (Inscopix, CA; **Figure 5A**). Our initial inquiry evaluated the impact of ELS on the magnitude of DA response following 40Hz VTA_BLA_ activation. We revealed that the VTA_BLA_-evoked DA response in CNT+ChR2 animals was reliably greater than baseline fluorescence, reaching significance within 1sec of VTA_BLA_ terminal activation (**Figures 5B-C, Stats Table 5**). However, ELS robustly decreased this response, quantitatively resembling CNT+eGFP animals (**Figures 5B-D, Stats Table 5**). Collectively, these data demonstrate that ELS reduces VTA_BLA_ DA signaling.

**Figure 5:**
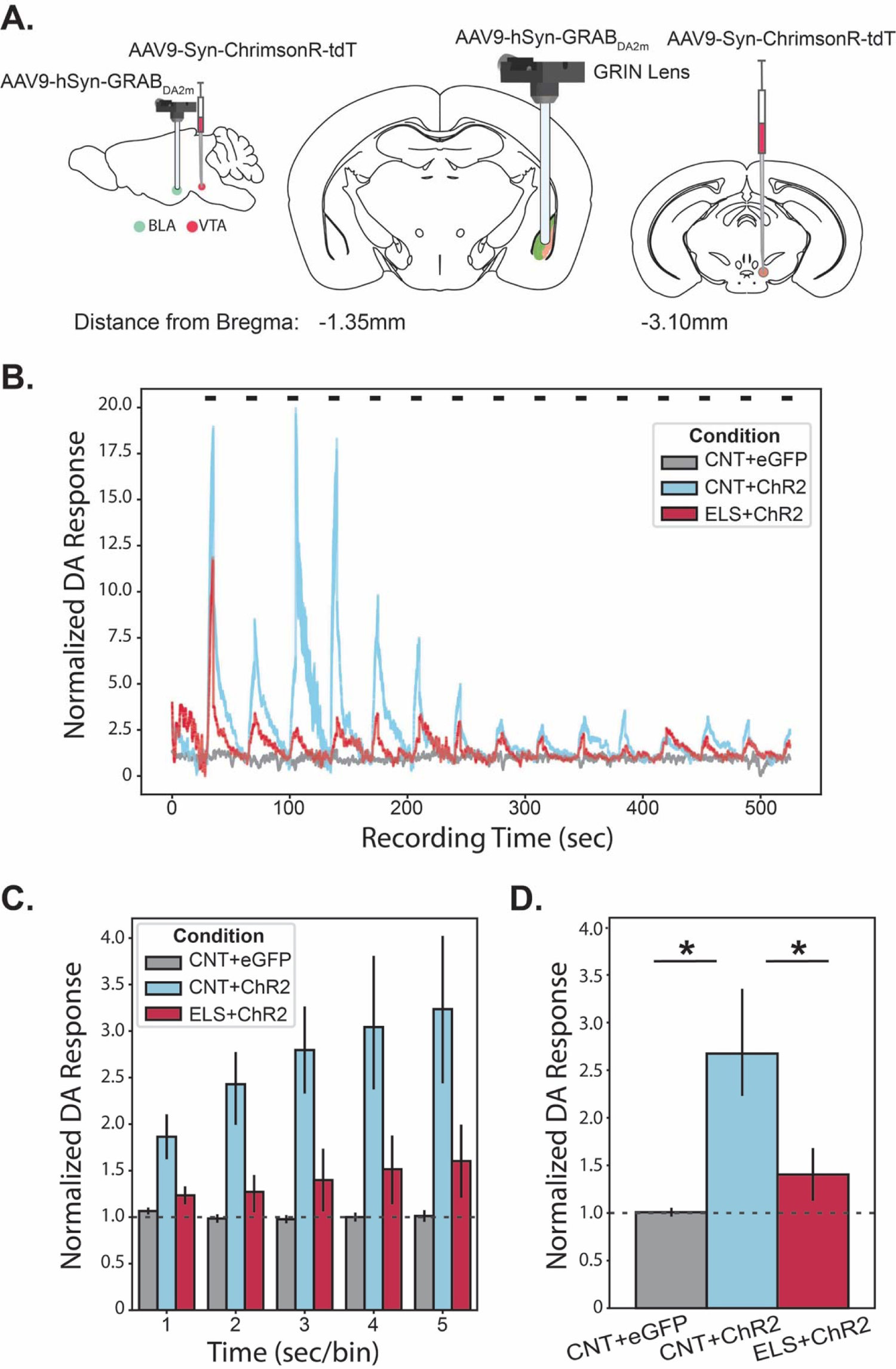
ELS reduces ELS reduces VTA_BLA_ DA response. (**A**) Schematic of sagittal and coronal views show AAV9-Syn-ChrimsonR-tDT or AAV9-hSyn-eGFP was injected into VTA, AAV9-hSyn-GRABDA2m injected into BLA, with GRIN lens implanted above BLA. (**B**) Representative traces show dopamine (DA) response across a recording window consisting of 15-5sec trials of 20mW red (620 +/- 30 nm) 40hz photoexcitation indicated by horizontal black bars; eGFP: grey, CNT: blue, ELS: red. (**C**) Grouped bar plot shows trial-averaged normalized DA response for each second of 5-sec stimulation for eGFP (n = 4), CNT (n = 4), and ELS (n = 5) animals. Dashed horizontal line indicates baseline response. (**D**) Summary plot shows session-averaged normalized DA response for YFP, CNT, and ELS animals. * *p* < 0.05. Error bars represent SEM.

Noting the impact of ELS on VTA_BLA_ DA signaling, we focused our attention on interrogating the impact of ELS on innate motivated behaviors, such as social interaction. We assessed the impact of ELS on social preference using a three-chamber social interaction task (3CST) which evaluated an animal’s proclivity to investigate either an inanimate object (fake mouse toy) or novel, female conspecific (female) with or without rhythmic VTA_BLA_ activation (**Figures 6A-D**). 40Hz stimulation was chosen for terminal activation because BLA gamma (30-80Hz) is known to be crucial for emotional/social processing [42, 54] and it is the frequency band where rearing conditions caused a significant difference in BLA-mPFC functional connectivity (**Figure 2**). CNT and ELS animals showed comparable interaction levels when presented with toys, with no effect of VTA_BLA_ photoexcitation (On) on interaction preference in comparison to trials in the absence of photoexcitation (Off; **Figures 6B** and **6E, Stats Table 6**). Conversely, when presented with the decision to interact with a toy or a female in the absence of VTA_BLA_ activation, animals robustly preferred to socialize, with ELS significantly enhancing this preference (**Figures 6D-E, Stats Table 6**) consistent with previous reports [55]. However, in trials where VTA_BLA_ activation occurred, ELS animals changed their preference, erring on the side of social avoidance, a behavior that was not exhibited in CNT animals (**Figures 6E-F, Stats Table 6**). These data suggest that the changes in VTA_BLA_ DA signaling impairs social interaction.

**Figure 6:**
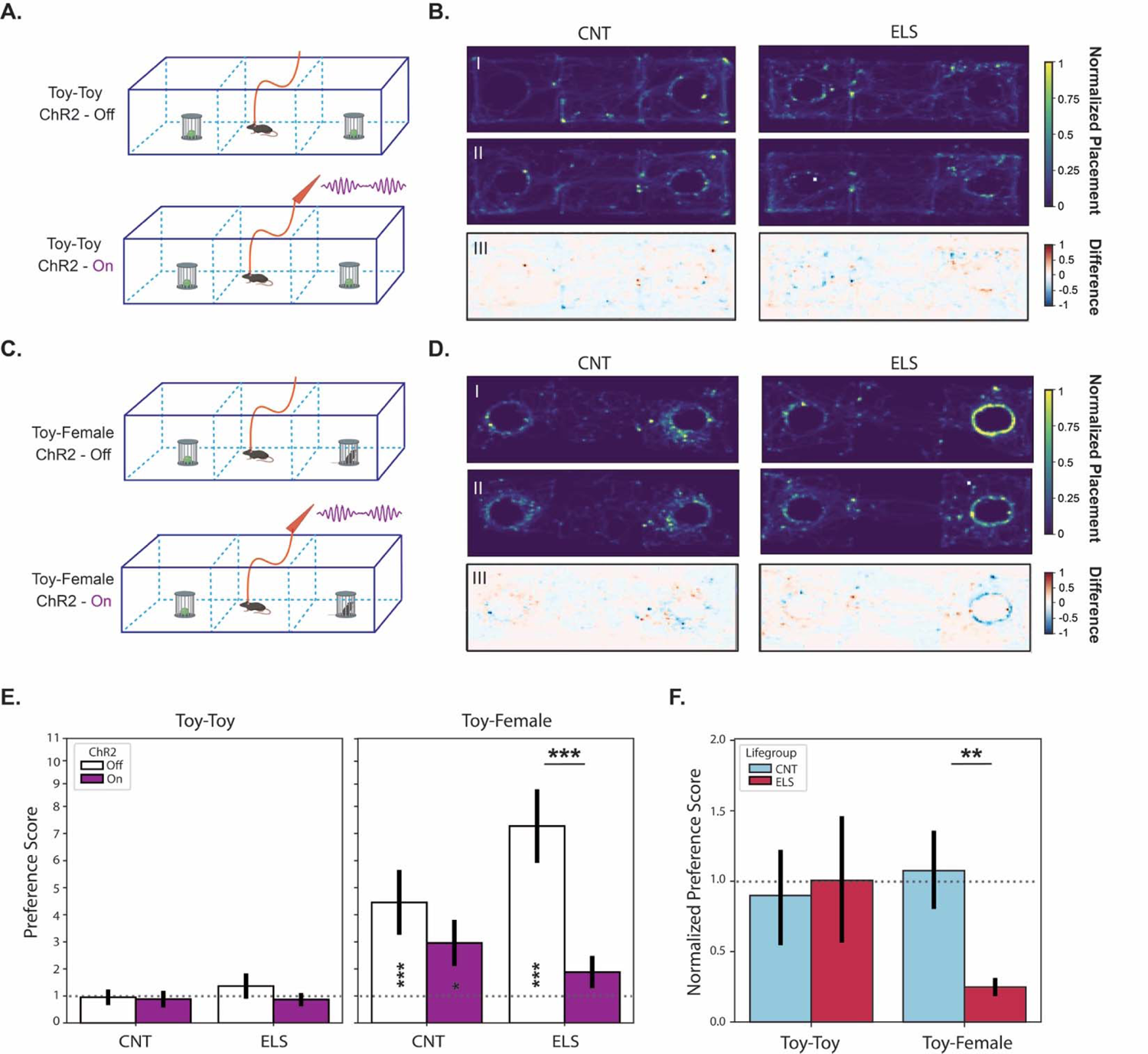
VTA_BLA_ activation in ELS reduces social interaction. (**A**) Schematic of 3-chamber social interaction task (3CST) during Toy-Toy sessions for habituation (ChR2-Off, Top) and testing (ChR2-On, Bottom) trials. (**B**) Representative heatmaps during 3CST show location for a CNT (Left) and ELS (Right) animal during habituation and testing trials, BI and BII respectively. Heat: Min-Max scaled duration across trial. BIII shows subtraction plots, with heat indicating the difference in time spent at each location between testing and habituation trials. (**C-D**) Same as (**A-B**) except interaction objects are a toy and novel, female conspecific (Toy-Female). (**E**) Grouped bar plots show interaction preference scores for CNT (n =7) and ELS (n=8) animals across habituation and testing trials for Toy-Toy (Left) and Toy-Female (Right) sessions; value of 1 is equivalent time spent in stimulation and opposite-side. Asterisks centered over trial: one side of chamber is significantly preferred. Asterisks centered across trials: preference scores during testing trials significantly differed from habituation trials. (**F**) Grouped bar plot shows normalized preference scores from (**E**) for CNT and ELS animals across Toy-Toy and Toy-Female sessions; value of 1: equivalent preference score between habituation and testing trials. * *p* < 0.05, ** *p* < 0.01, *** *p* < 0.001. Error bars represent SEM.

To directly test whether impaired functional connectivity between the VTA and BLA following ELS contributes to altered social interaction, we silenced VTA_BLA_ signaling and examined the impact on social preference. In an adaptation of the experimental protocol detailed in **Figure 1A**, we infused a red-light sensitive chloride pump-encoding viral vector (AAV-GFP-JAWS; Jaws) into VTA to promote VTA_BLA_ terminus expression aimed at inhibiting VTA-mediated activity (**Figure 7A**). Given that we saw an ELS-specific reduction in VTA_BLA_ signaling, we posited that inhibiting VTA_BLA_ information in CNT animals would result in a similar social avoidance behavior exhibited by ELS animals. CNT+Jaws mice were then subjected to the same 3CST detailed in **Figure 6**, with the exception that during testing trials, VTA_BLA_ signaling was inhibited (**Figures 7B-C**). Interaction levels were comparable between habituation and testing trials when CNT+Jaws mice were presented with toys. Meanwhile, when presented with a female conspecific, these animals significantly increased their social interaction; a preference that was significantly decreased during VTA_BLA_ inhibition (**Figure 7D, Stats Table 7**). When we directly compare the results between the VTA_BLA_ excitation and inhibition experiments, our hypothesis is borne out: VTA_BLA_ inhibition causes a reduction in socializing in CNT animals, a VTA_BLA_-induced phenotype exhibited following ELS, which is not observed in VTA_BLA_-stimulated CNT animals (**Figure 7E**).

**Figure 7:**
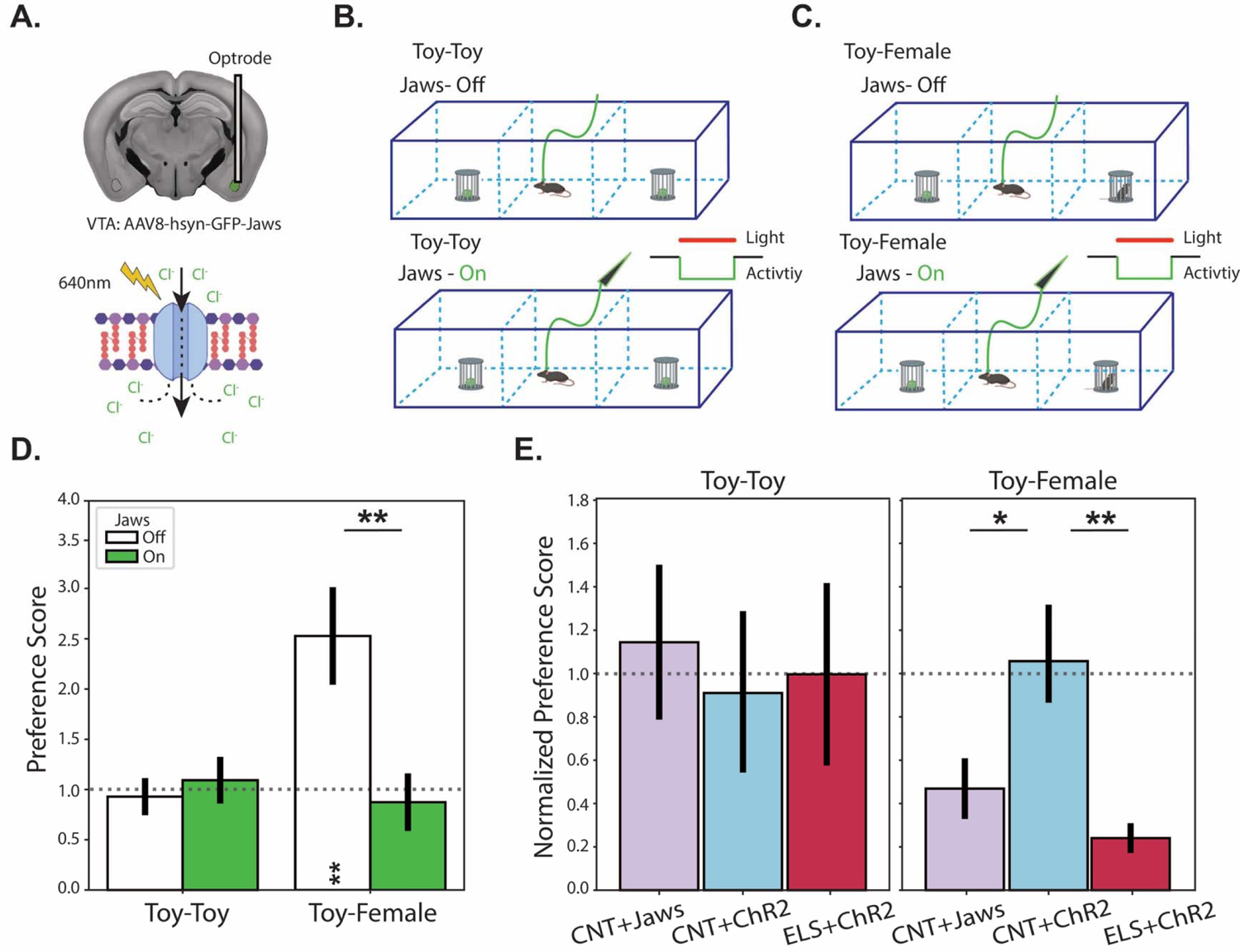
VTA_BLA_ inhibition mimics social deficit observed following ELS + VTA_BLA_ excitation. (**A**) Top: Schematic of coronal slice showing AAV8-hsyn-GFP-Jaws was injected into VTA with optrode implanted into BLA. Bottom: Schematic of inhibitory mechanism of red-shifted Jaws acting on anion-gated membrane following 640nm photostimulation. (**B**) Schematic of 3CST during Toy-Toy sessions for habituation (Jaws-Off, Top) and testing (Jaws-On, Bottom) trials. (**C**) Schematic showing same experimental set-up as in (**B**) with the exception being interaction objects are a toy and novel, female conspecific (Toy-Female). (**D**) Grouped bar plots show interaction preference scores for Toy-Toy and Toy-Female sessions habituation and testing; value of 1 is equivalent time spent in stimulation and opposite-side. Significance stars centered over trial: one side of chamber is significantly preferred. Significance stars centered across trials: preference scores during testing trials significantly differed from habituation trials. (**E**) Grouped bar plot shows normalized preference scores from (**D**) for CNT+Jaws animals (n = 5) as well as CNT+ChR2 and ELS+ChR2 animals (from **Fig 6**) across Toy-Toy and Toy-Female sessions; value of 1: equivalent preference score between habituation and testing trials. * *p* < 0.05. ** *p* < 0.01.

## Discussion

The results of this study advance our understanding of the mechanisms through which early life stress affects brain circuitry and behaviors in mice. The findings highlight the importance of VTA_BLA_ signaling in coordinating BLA-mPFC activity and influencing social preference, with ELS disrupting this coordinated signaling and leading to altered social behavior. Specifically, we observed that rhythmic activation of VTA_BLA_ terminals entrained BLA network activity, elevated BLA-mPFC coherence, increased BLA dopamine levels (DA), and had no effect on social behavior in control mice. In contrast, following ELS, we observed weakened VTA-mediated BLA and mPFC entrainment, diminished BLA-mPFC coherence, reduction of VTA_BLA_ DA terminals together with reduced BLA DA levels. Interestingly, ELS mice also exhibited activity-induced social avoidance. Given the pervasiveness of ACEs and their well-established comorbidity with psychiatric disorders [5–7], the elucidation of the mechanism behind such vulnerabilities remains paramount. Our study highlights the pivotal role of the MCL system, more specifically the interaction between VTA and BLA, in mediating these behavioral outcomes.

Maternal neglect has been robustly associated with cognitive impairments and increased susceptibility to psychiatric disorders [1–3]. Our results support this by demonstrating that ELS can modulate the neurophysiological pathways implicated in several stress-linked behaviors impacting social reward. However, while the involvement of the MCL system (particularly the MCL DA system) in stress vulnerability has been documented [8, 9, 12], our study provides a breakthrough by elucidating the impact of VTA DA on BLA and mPFC network dynamics, bringing to light how ELS affects their functional connectivity which is known to influence behavioral states. These results reveal that ELS affects the quantity of BLA-targeted VTA DA neurons and thus alters DA release, which could impair BLA-mPFC information flow. Given that reduced DA tone in the prefrontal cortex (PFC) has been associated with impaired cognitive function and increased susceptibility to stress (for a review see [56]), and that altered connectivity between the PFC and the amygdala leads to heightened emotional reactivity and impaired regulation [57], we propose that ELS alters social behaviors by impacting DA circuitry through the BLA. Our findings underscore the critical interplay between these regions, especially in the context of ELS; disrupting the BLA-mPFC network has profound behavioral consequences, notably social avoidance, thereby offering potential pathways for therapeutic interventions.

There is precedence in the literature emphasizing the critical role of the BLA in processing salient stimuli [40, 41]. Our results further suggest that the integrity of the VTA_BLA_ pathway might be vital for maintaining this capability, with disruptions leading to maladaptive behaviors. The specific mechanisms uncovered, namely a reduction in BLA-directed DA afferents from the VTA and diminished BLA DA sensitivity, provide deeper insight into potential therapeutic targets. These findings are especially significant in light of the fact that DA signaling in the BLA, and its downstream pathways, plays a pivotal role in motivation and reward processing, features that are frequently altered in individuals with psychiatric disorders.

While some of these findings may initially appear to be in conflict – namely the reduction in DA signaling and BLA-mPFC coherence that occurs concurrent with an increase in social avoidance observed in ELS mice – we posit that social avoidance is governed by a delicate balance of interconnected neural elements. We hypothesize that several components (likely working in concert) explain how ELS shapes VTA-BLA dynamics in a nuanced relationship requiring VTA_BLA_ DA input, rhythmic modulation, and sensory input(s) that are intrinsically salient. Specifically, the observed reduction of VTA_BLA_ DA projections following ELS has the propensity to alter communication between BLA and mPFC which is well-established to regulate social behaviors [58, 59]. Further, the rhythmic activity driven by VTA likely acts as a trigger, promoting or inhibiting inter-region information transfer [60]. This could explain why VTA_BLA_ activation in control animals does not disrupt the existing balance of signal transmission, resulting in no effect on social preference; only when the VTA_BLA_ DA circuit is inhibited does the dopamine response change to maladaptive.

Further, it has been shown that ELS can induce structural and functional changes in brain, regions related to stress, emotion, and social behaviors [61, 62], specifically altering synaptic plasticity, dendritic morphology, and the density of neurotransmitter receptors within these regions ([63], see also **Figure 4**). Studies have also shown that ELS-induced anhedonia (loss of pleasure capacity) has been linked to aberrant signaling interactions across the MCL circuit [2, 64] (for review see [65]). To maintain homeostasis, therefore, one likely explanation for our results is that ELS animals develop compensatory mechanisms, such as strengthening synaptic connections (e.g., synaptic efficacy), remapping of certain neural pathways, or balancing of excitatory/inhibitory signaling. These mechanisms together, or in and of themselves, could lead to differential behavioral outcomes, particularly when this MCL network is challenged (e.g., VTA_BLA_ excitation or inhibition). Accordingly, we propose that the MCL system is targeted during ELS and that the observed alteration in social motivation is substantiated in VTA DA driving (or failing to drive) oscillations across the BLA and mPFC which govern interaction behaviors.

Our study offers a nuanced understanding of the role of VTA-BLA connectivity in mediating social motivation and how ELS can impair this functionality. Our findings pave the way for future research on potential therapeutic interventions, emphasizing the restoration or modulation of this pathway as a possible strategy. Further investigations into the molecular and cellular mechanisms underlying these changes could provide valuable insights into the pathophysiology of social deficits in psychiatric conditions, such as major depressive disorder and social anxiety disorder. While further research is undeniably required, especially in understanding the broader implications of these findings, our study constitutes a significant step in understanding the neural underpinnings of stress vulnerability.

## Methods

### Animals

C57BL/6J mice were housed at Tufts University School of Medicine in a temperature and humidity-controlled environment, maintained on a 12-h light/dark cycle (lights on at 7 a.m.) with ad libitum access to food and water. All animal procedures were handled according to the protocols approved by the Tufts University Institutional Animal Care and Use Committee (IACUC).

### Early Life Stress Paradigm

Pregnant C57BL/6J dams were purchased directly from Jackson Laboratory (Bar Harbor, ME) and were delivered and singly housed on day 16 of gestation (E16) at Tufts University School of Medicine. These cages were checked daily for litter births, at which time litters were randomly assigned to standard-rearing (control; CNT) or maternal separation (early life stress, ELS) groups. ELS began on postnatal day 1 (PND1) and consisted of the removal and placement of the dam into a clean cage, out of the sight of her litter, for 3 hours (beginning at 9AM) for 5 days per week from PND1-21. Offspring were weaned on PND21.

### Stereotaxic Surgery

Adult (>PND60) male CNT and ELS mice were anesthetized by intraperitoneal injection (IP) of a ketamine/xylazine cocktail (90 – 120 mg/Kg and 5 – 10 mg/Kg, respectively). Before the onset of the procedure, sustained release-buprenorphine (0.5 – 1 mg/Kg) was administered subcutaneously as post-operative analgesia. The head of the mouse was shaved, antiseptic solution was applied, and an incision on the scalp was made to expose the skull. For viral injections, mice were unilaterally injected with a virus targeted at either the basolateral amygdala (BLA) or ventral tegmental area (VTA; see **Methods Table 1** for details) using a pulled glass pipette at a flow rate of 100LJnL per minute using positive pressure from a 10 mL syringe. All viruses were purchased from a Addgene. Post Hoc imaging was performed to validate targeted viral expression. For the voltage-clamp and retrograde labeling experiments (**Figures 3 and 4**), brains were collected 2.5 weeks post-viral injection.

**Methods Table 1.**

| Figure | Virus | Titer<br>(vg/mL) | Volume<br>(nL) | Dilution | Region | From bregma<br>(A/P, M/L, D/V) |
| --- | --- | --- | --- | --- | --- | --- |
| **Figure 1, 2, 5, 6, 7** | AAV9-syn-ChrimsonR-tdT; ChR2 | $1 \times 10^{13}$ | 300 | NA | VTA | -3.10, -0.53, -4.50 |
| **Figure 4** | rAAV-GCamp6f-tdT | $1 \times 7^{12}$ | 200 | NA | BLA | -1.35, -3.20, -5.10 |
| **Figure 5** | AAV9-hSyn-GRAB<sub>DA2m</sub> | $1 \times 10^{13}$ | 600 | 1:2 | BLA | -1.35, -2.90, -4.70 |
| **Figure 5** | AAV9-hSyn-eGFP | $1 \times 7^{12}$ | 300 | NA | VTA | -3.10, -0.53, -4.50 |
| **Figure 7** | AAV-hsyn-Jaws-KGC-GFP-ER2 | $1 \times 10^{13}$ | 300 | NA | VTA | -3.10, -0.53, -4.50 |

### LFP recordings

LFP recordings were performed in awake, adult C57BL/6J CNT and ELS mice, using custom-designed headmounts (Pinnacle 8201-C). These headmounts consisted of two depth electrodes (PFA-coated stainless-steel wire, A-M systems) targeted at the medial prefrontal cortex (mPFC; from bregma: AP: +1.75; ML: −0.30; DV: −2.0mm) and BLA (from bregma: AP: −1.35; ML: −3.20; DV: −5.10mm). The headmounts were affixed to the skull using stainless steel screws as ground and reference following 2.5 weeks from viral injection. The LFP data were band-pass filtered (1-250 Hz, Chebyshev Type II filter) and spectral analysis was performed in Python using custom-made scripts utilizing the fast Fourier transform. Specifically, spectral analysis of the LFP/EEG signal was performed using the Short Time Fourier Transform (STFT) with a 5 second Hann window and 50% overlap as done previously[17]. The power line noise (59 - 61 Hz) was removed and values were filled using nearest interpolation. Outliers in each spectrogram were identified using a two-stage process. Firstly, a time-series was obtained from the mean power across frequencies of each spectrogram. Large deviations, defined as those greater than the mean plus 4 times the standard deviation, were replaced with the median of the non-outlying data. Then, a sliding-window method was applied to detect more subtle outliers based on local context, using 5x the Median Absolute Deviation (MAD) of each 5-minute window. Resulting outliers were removed and replaced with forward fill interpolation of the nearest values. All resulting time-series, obtained from the mean power across frequencies, were manually verified.

For optogenetic experiments, CNT and ELS mice expressing either ChR2 or Jaws (**Methods Table 1**), were implanted with an in-house modified version of the described headmounts, optrode, which consisted of the addition of an optic fiber (200 μm, 0.22 NA; ThorLabs) attached to the depth electrode aimed at BLA. Photo-stimulation was performed with a red light delivered through the optic fiber coupled to a red laser (640 nm, max power=500 mW, Laserglow technologies). The light intensity setting used for all optogenetic experiments in awake mice was measured at 30-40 mW from the end of the opto-cannula (Thorlabs, Inc.; 200 μm, 0.39 NA). Sine waveforms for photo-stimulation were generated through LabChart (ADInstruments). The LFP data were band-pass filtered between 3-300 Hz.

For pulsed photo-stimulation experiments (5, 10, 20, and 40Hz; **Figures 1 and 2**), stimulations were pseudo-randomized, where each stimulation frequency was repeated 3 times, and post-hoc analyses revealed no presentation order effect on power elevation (**Supp.** Fig 1E**/F**). The average response was used for further quantification. The power spectral densities for optogenetic experiments were obtained in 1 second bins (50 % overlap) using custom-made Python scripts. For LFP power ratio (**Fig. 1C/D** and **Supp.** Fig 1E**/F**), quantification of the power of the oscillations was obtained around +/- 1 Hz of each stimulation frequency during (0 − 120 sec.) and before stimulation (0 − 30 sec.). Only stimuli that produced phase-locked LFP responses +/- 2 standard deviations from group average, were included. The Continuous Morse Wavelet Transform [66] was used to visualize opto-spectrograms. For LFP coherence analyses (**Figure 2**), BLA-mPFC LFP synchronization was quantified using the Phase Locking Value (PLV, see [67]), obtained around bandpass-filtered +/- 1 Hz of each stimulation frequency, using custom-made Python scripts. These values were normalized across frequencies as a ratio between stimulation period and baseline period, as done for power-ratio analyses.

### Current-clamp recordings

Brains from adult CNT and ELS mice were extracted in ice cold (0-4 °C) aCSF solution containing (in mM) 126 NaCl, 10 glucose, 2 MgCl2, 2 CaCl2, 2.5 KCl, 1.25 NaHCO3, 1.5 Na-pyruvate, 1 L-glutamine (300 – 310 mOsm) and 3 mM kynurenic acid. Coronal brain slices (300 μm) were obtained and transferred to a submerged chamber in a warm (∼ 34° C) normal aCSF for at least 1 hour prior to recordings. Slices were then transferred to a submerged recording chamber. Borosilicate glass electrodes were pulled with a resistance of ∼3-5 MΩ (DMZ Universal Puller). The intracellular recording solution contained (in mM) 130 mM potassium gluconate, 10 mM KCl, 4 mM NaCl, 10 mM HEPES, 0.1 mM EGTA, 2 mM Mg-ATP, and 0.3 mM Na-GTP (pH 7.25; 280–290 mOsm). Data were acquired at 10 KHz and were low pass filtered at 3 KHz (AD Instruments). Recordings were performed in principal neurons in the BLA identified based on morphology. Current clamp recordings were performed to generate input-output curves in response to a series of current injections from 0-150pA in 10pA steps. Intrinsic electrophysiological properties, including resting membrane potential, input resistance, and impedance. The electrophysiological properties were measured before and after a 5 min bath application of SCH 23390 (5µM). Cells were eliminated from analysis if the series resistance was greater than 20 MΩ or changed >20% over the course of the experiment.

Action potentials (APs) were detected using SciPy’s *find_peaks* function (prominence=50 mV, wlen=100 ms, distance=1 ms). The IO slope was calculated using a first-degree least squares polynomial fit (NumPy’s *polyfit*, n=1) between rheobase to 80% of max firing rate. For the IO slope, cells were included only if they fired for at least 3 steps from rheobase to 80% of max APs (at least 3 points for line fit). For waveform analysis, only a maximum of 3 APs were collected from up to the first 3 current steps above rheobase to ensure that the afterhyperpolarization (AHP) was maximally preserved ([68]; **Supp2**). To calculate waveform properties, APs were interpolated (SciPy’s *interp1d*, interpolation factor=10, type=cubic). The AP threshold was detected as 1.5 V/s, the AP amplitude was defined as the difference from the AP peak – threshold. The AHP amplitude was defined as the absolute difference between the AHP peak – AP threshold. AP peak-to-trough was calculated as the time between AHP peak to AP peak. The rise time was calculated as the time between AP peak to AP threshold. For the analysis of the effects of D1R antagonist (SCH 23390) on extracted properties, only cells that exhibited APs before and after drug application were included. For the extraction of Vm, cells that did not exhibit a stable baseline were removed upon manual inspection. The signal was first downsampled to 10 Hz with an anti-aliasing filter (Scipy’s *decimate*). Outliers were detected using a sliding-window method using 4x the Median Absolute Deviation (MAD) of a 90 second window. Outlier regions were then defined as merged outliers within 2 seconds and further expansion of the merged margins by 2 seconds on each side. Outlier regions replaced with linear interpolation (Pandas *interpolate*). Finally, the remaining outlier free time series was smoothed using a median filter (0.2 seconds, *Scipy’s medfilt*),

### Immunohistochemistry

Immunohistochemistry was performed on coronal sections (50μm thick) which were rinsed with PBS and then blocked with buffer (PBS, 1% bovine serum albumin (BSA), 5% fetal calf serum (FCS), and 0.2% Triton X-100). This was followed by overnight incubation with primary antibody (see below) and eventually conjugated to a fluorophore with the secondary antibody (1:500, ThermoFisher, Alexa Fluor). The sections were incubated in primary antibody for 24 h at 4 °C, washed 2X with PBS, and then incubated, covered, with secondary antibody for 2 h at room temperature. After washing 3X with PBS the sections were incubated with DAPI (1:20,000; catalog no. D1306, Invitrogen) and the sections were mounted on Superfrost slides (Fisher Scientific; catalog no.12-550-15) and cover slipped with Prolong Gold Antifade (catalog no. P36930; Invitrogen).

**D1-receptor expression**: Primary; chicken anti-DA receptor D1a (1:600; Millipore MAB5290)

**TH expression:** Primary; chicken anti-TH primary antibody (1:1000; Abcam, AB76442)

A wide-field epifluorescence microscope (Keyence BZ-X700) was used to acquire images of mounted sections. Images for D1-receptor expression were taken at 20x, while images for TH expression were taken at 10x. Images were stitched together using Keyence software. Cell counts, receptor expression/density, and colocalization were performed using CellProfiler software (version 4.2.5, Broad Institute, Cambridge, MA; www.cellprofiler.org). Modified versions of CellProfiler pipelines (pipelines and detailed instruction can be obtained from https://cellprofiler.org/examples) were adapted to our specificities, where dimensions of cells and fluorescent levels were changed to reliably quantify images. Given our 50μm slice thickness, and that VTA dopaminergic neurons are <30μm (ranging between 10-25μm; [69], for review see [70]), each slice was treated as its own observation for statistical purposes.

### Recording in vivo dopamine responses in BLA

Biosensing of dopamine binding was performed in awake, adult CNT and ELS animals, using the GRAB-DA sensor and the nVoke miniaturized microscope (an integrated imaging and optogenetics system, 455-nm blue GCaMP excitation LED, 590-nm amber optogenetic LED, Inscopix). Following injection of ChR2 in VTA and GRAB_DA2m_ in BLA (**Methods Table 1**), an Inscopix lens affixed to a baseplate (Inscopix, 1050-004413) was placed in the BLA. Mice were allowed to recover for 7 weeks prior to experimentation to allow for optimal viral expression and *in vivo* imaging. On the recording day, each animal was removed from their home cage, placed in a testing cage (a cage identical to home-cage, containing fresh bedding/nesting), and had the miniature integrated microscope system (nVoke HD 2.0; Inscopix) attached to their baseplate. Each animal was given 10 min to habituate to the testing cage, at which time images were acquired using data acquisition software (ver. 2.0.0; Inscopix) at 20 frames per sec, 20% of LED power, and a gain kept between (2 and 5, varying based on clarity of field of view; FOV). Images were acquired across a 10 min session consisting of 15 trials of 40hz, 5-second pulse trains (5 ms width) of 620-nm LED (20mW) light initiated every 30 seconds.

Acquired imaging data were down-sampled (1/2 spatial binning), preprocessed, motion corrected, DA transients of ROIs (3 concentric circles as detailed in Inscopix Processing Guidelines: https://iqlearning.inscopix.com/best-practices/applications/nt-imaging/single-color/processing) were extracted, and a change in fluorescence (ΔF/F) was computed using Inscopix Data Processing Software (ver. 1.9.2; Inscopix). All extracted traces were then subjected to a series of custom-built Python scripts which: quadratically-detrended the DA signals (to account for photo-bleaching across a session; application of a second-degree polynomial fit as seen in [71, 72]), normalized each ROI to the mean baseline (pre-stimulation) periods, and then parsed these signals into individual trials for analyses. Given that the DA response is intrinsically linked to the available DA in the region of recording, all statistical tests were computed using the trials as observations, thus accounting for the dependent nature of DA depletion across a session.

### Three-chamber social interaction task (3CST)

We recorded behavior from adult CNT and ELS mice during a 3CST following 3 weeks post-viral infusion in a Plexiglas rectangular box (71 cm X 30 cm X 35 cm), made up of three identically shaped compartments, without a top. The compartments were separated by a transparent piece of Plexiglas, with cut-outs to allow for free movement between each compartment. The left and right compartments each had an inverted wire cup (diameter 8 cm) placed in the center of them, which had a designated depressed (1/4 inch lower) cylindrical section to ensure wire cups remained in place throughout testing. The apparatus and wire cups were thoroughly cleaned with 70% ethanol between each trial.

The 3CST consisted of two sessions (Toy-Toy and Toy-Female), each made up of a habituation and testing trial. The testing protocols were identical in structure made up of 4 trials, each with the inverted cups holding either a toy or a novel, female conspecific: mice explored the arena and toys (Toy-Toy session; habituation) for 10min, they repeated this with the exception that one of the sides (testing; pseudo-randomly chosen) triggered photostimulation upon physical entrance into the interaction zone (3cm space directly extending beyond the wire cup), they begin a new habituation trial where they can explore and interact with a toy or a female mouse (Toy-Female; habituation), and finally they explore the arena with the same female mouse, triggering photostimulation upon entrance into the interaction zone around her wire cup (testing). During interaction-zone triggered stimulation, mice received VTA_BLA_ 30-40 mW photo-stimulation through the optic fiber coupled to a red laser (640 nm) at 40Hz (**Methods Table 2**).

**Methods Table 2:** 3CST Sample Protocol. Stim., Stimulation; R, Right; L, Left.

| Animal | Session | Trial | Stim. | Stim. Side | Detail |
| --- | --- | --- | --- | --- | --- |
| Animal 1 | Toy-Toy | Habituation | Off | -- | Toy inside cage on both sides of area |
|  |  | Testing | On | L | Toy inside cage on both sides of area |
|  | Toy-Female | Habituation | Off | -- | Toy inside L cage, Female inside R cage |
|  |  | Testing | On | R | Toy inside L cage, Female inside R cage |
| Animal 2 | Toy-Female | Habituation | Off | -- | Toy inside R cage, Female inside L cage |
|  |  | Testing | On | L | Toy inside R cage, Female inside L cage |
|  | Toy-Toy | Habituation | Off | -- | Toy inside cage on both sides of area |
|  |  | Testing | On | R | Toy inside cage on both sides of area |
| Methods Table 2: 3CST Sample Protocol. Stim., Stimulation; R, Right; L, Left. |  |  |  |  |  |

The overall activity of the test mouse in the 3CST was automatically recorded with the EthoVision XT video tracking system (Noldus, Netherland). The amount of time that test mice spent in each chamber and the immediate vicinity (3 cm) of the cages (termed as “interaction zone”) was measured. A custom-build Arduino script was used to receive inputs from EthoVision XT, which detailed when an animal was in the interaction zone, causing the 640nm laser to send the 40hz pulse trains (described above) indefinitely, until the animal left the interaction zone. Social preference scores (*PS*) were calculated as the percentage of time spent in the stimulation side (*T_S_*) out of the total time spent in either left or right side of the area (*T_S_ + T_NS_*; see Equation 1). Normalized preference scores (*NS*) were calculated as the ratio between the preference score observed during the testing trial (*PS_T_*) out and the preference score observed during the habituation trial (*PS_H_*; see Equation 2). Here, time (*T*) is the cumulative seconds that the nose of the test mouse was detected in the interaction zone (which ensures actual interaction between test mouse and caged object; toy or female mouse). In this manner, we were able to unbiasedly quantify interaction preference, whereby a PS value greater than 1 indicates more time spent interacting on the stimulation side. An NS value greater than 1 indicates the 640nm stimulation (testing trials) increased this interaction time, in comparison to non-stimulated (habituation) trials.

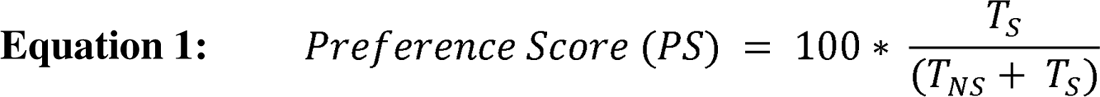

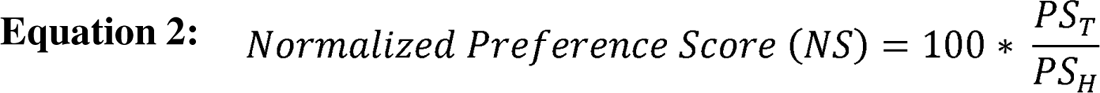

Lastly, to minimize inter-animal variability across CNT/ELS groups and viral manipulations (excitatory and inhibitory), female conspecifics in estrous (most receptive to mating) using vaginal lavage and microscopy according to [73] on the day of testing. Groups were counterbalanced for order of light stimulation as well as side assigned as the social zone (see **Methods Table 2** for details). A 3way mixed-ANOVA (lifegroup*stimulation side*session) was performed and revealed no significant difference (*p* >0.05), revealing that the session presentation order, nor side the stimulation occurred on, affected CNT and ELS animals differently.

### Quantification and statistical analysis

All statistical analyses were performed using Python (Python Software Foundation). Multi-group tests were tested with ANOVAs followed by post-hoc multiple comparisons. A Greenhouse-Geisser correction was applied where appropriate and when assumption of sphericity was violated. For electrophysiological parameters between two groups Mann-Whitney U tests were run with Benjamini/Hochberg procedure to control for False Discovery Rate (statsmodels). For comparison of ELS and CNT, appropriate one-way tests were applied, chosen based on distribution of data and test assumptions. The statistical package Pingouin was used for all tests, except for correlation analyses and patch-clamp electrophysiological comparisons. All tests and results are reported for each comparison, named in accordance with the figure the results detail. A p-value <0.05 was considered statistically significant.

## Acknowledgements

This work is supported by National Institutes of Health (Grant Nos. R01AA026256, R01NS105628, R01NS102937, R01MH128235, P50MH122379). We would also like to thank Drs. Morgana Favero and Sinem Erisken for their assistance in implementing Inscopix imaging in the laboratory.

## Author Contributions

B.S. and J.M. contributed to the conceptualization of the project; B.S., E.T., G.S., and J.M contributed to investigation; B.S., P.A., and G.W. contributed to analyses; B.S. and J.M. contributed to writing—original draft; B.S., P.A., E.T., G.S., G.W. and J.M. contributed to writing—review and editing; J.M. obtained the funding for the project.

## Materials and Correspondence

Further data supporting the findings are available upon reasonable request to the corresponding author.

---

**Supplemental Figure 1:**
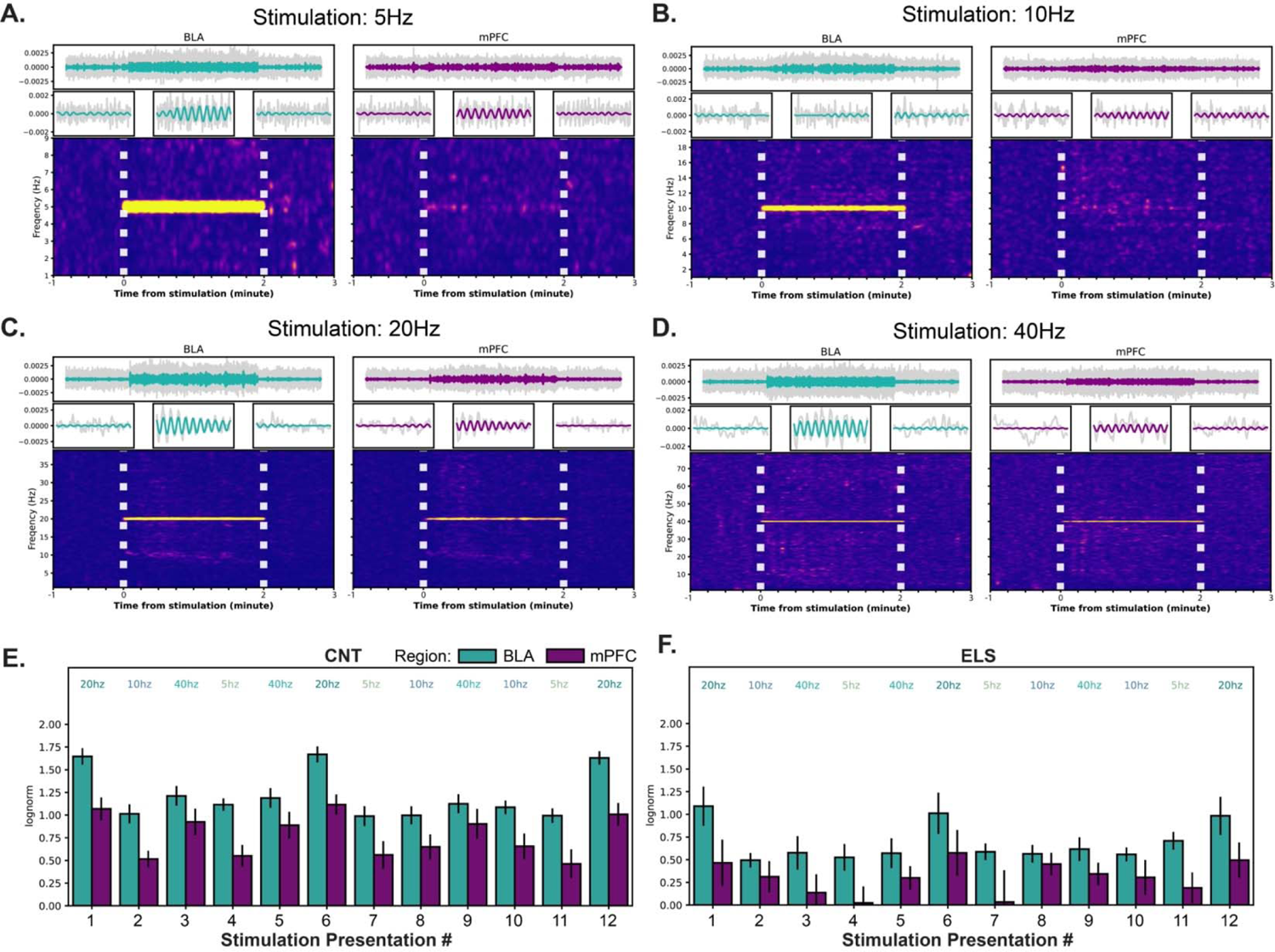
VTA_BLA_ robustly drives BLA and mPFC network activity. Representative traces and wavelet transformations for a CNT animal during a (**A**) 5Hz, (**B**) 10Hz, (**C**) 20Hz, and (**D**) 40Hz trial. Top: Stimulated frequency LFP trace atop filtered (2-100Hz) LFP traces (grey) from BLA (left) and mPFC (right) for; Middle: Zoomed in portions from above traces show 5 periods before, during, and after 10mW 640nm photoexcitation; Bottom: Wavelet transformation from respective traces; horizontal white lines indicate onset/offset of 640nm laser. (**E**) Grouped bar plot, organized by stimulation order, shows LFP power ratio between pulsed stimulation and baseline periods where power was measured at each stimulation frequency for CNT (blue, n = 7) animals. (**F**) Same as (**E**) with the exception that analyses performed on ELS (red, n = 8) animals.

**Supplemental Figure 2:**
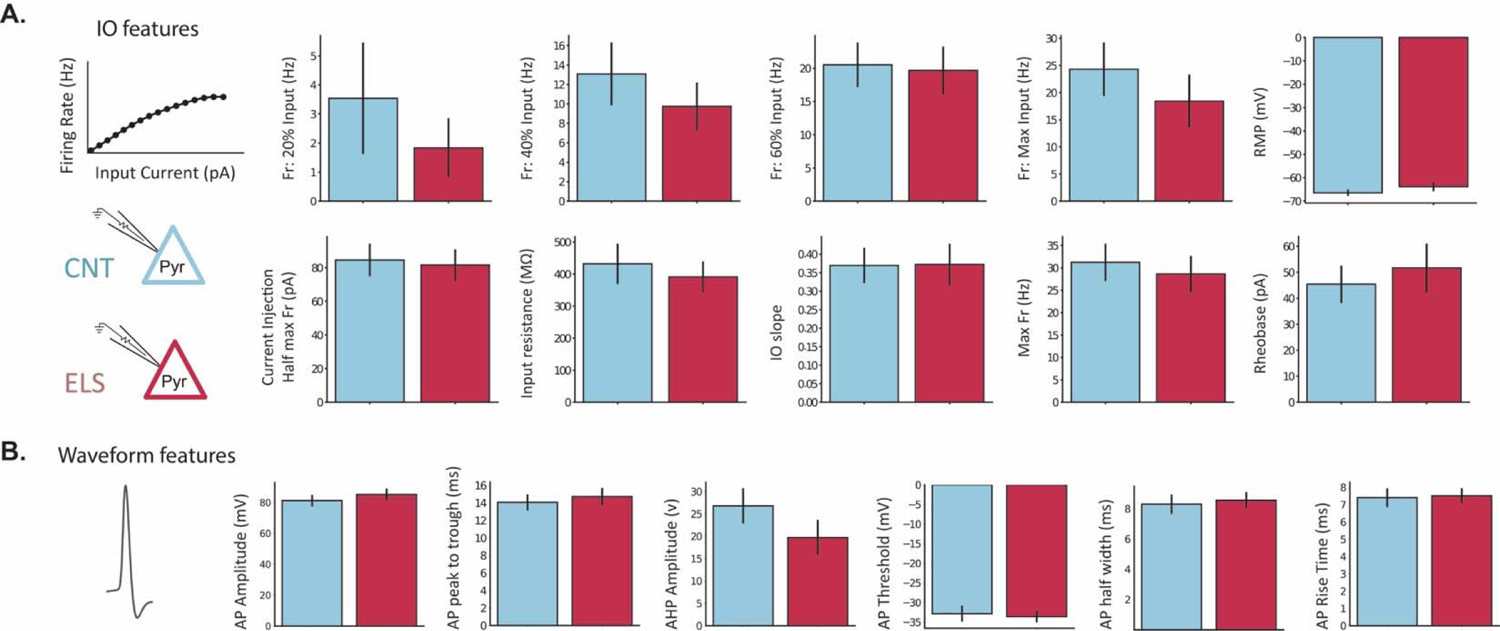
Effect of ELS on BLA principal cell properties. (**A**) Bar plots for input-output properties. (**B**) Bar plots for action potential waveform properties. Error bars represent SEM.

**Supplemental Figure 3:**
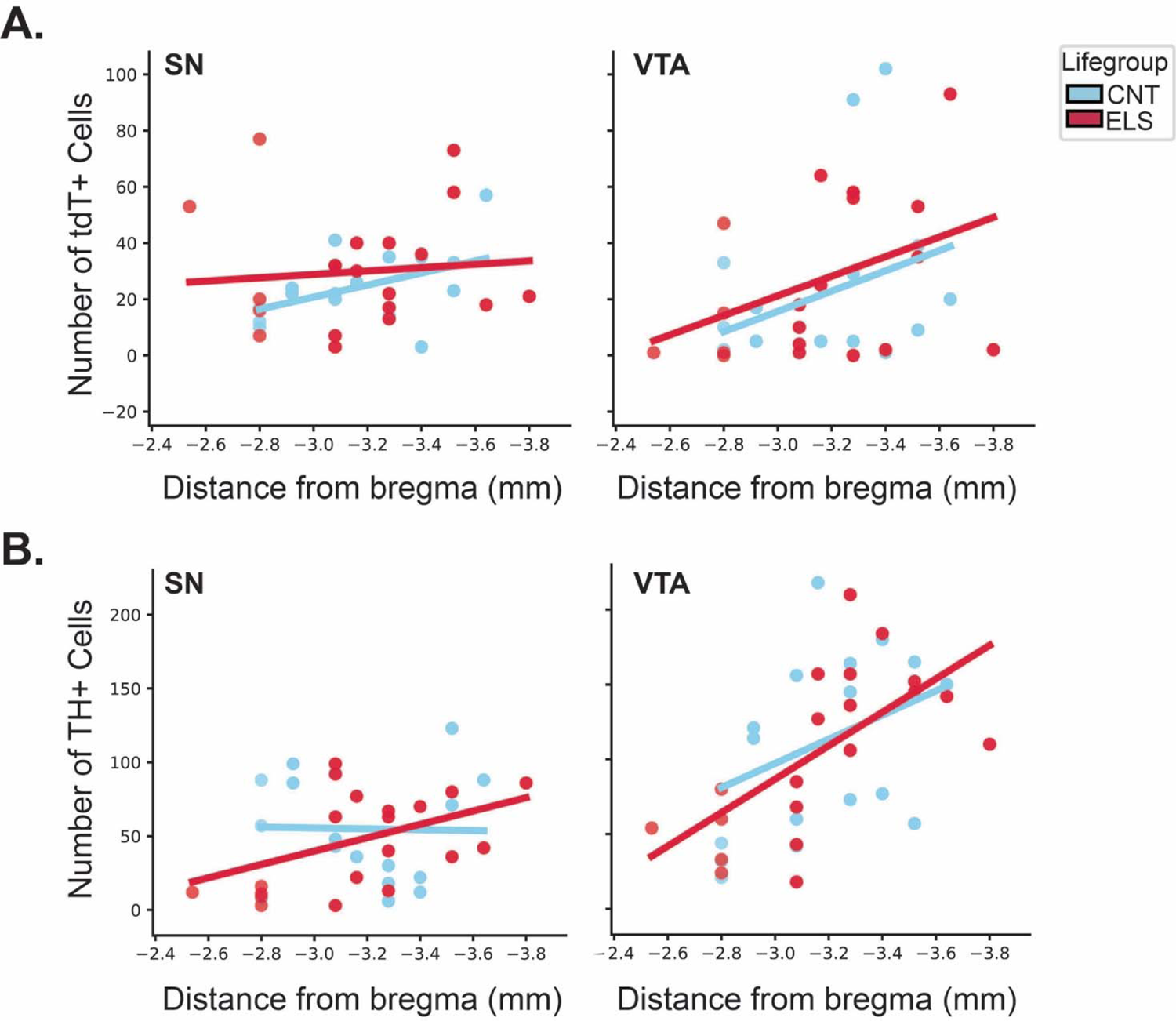
Localization and number of tdT+ and TH+ neurons in SN and VTA. (**A**) Cell counts for BLA-projecting VTA neurons (tdT+) given per 50-μm-thick coronal section as mean per hemisphere/slice in CNT (N = 5 animals, n = 6 slice per animal) and ELS (N = 4 animals, n = 6 slice per animal) animals at different levels from bregma along the rostro-caudal axis; Left: SN, Right: VTA. **(B**) Same as (**A**) with the exception being analysis of TH+ cells. Regression lines in each plot correspond to Pearson correlation coefficients presented in Stats Table 4.

**Stats Table 1.**
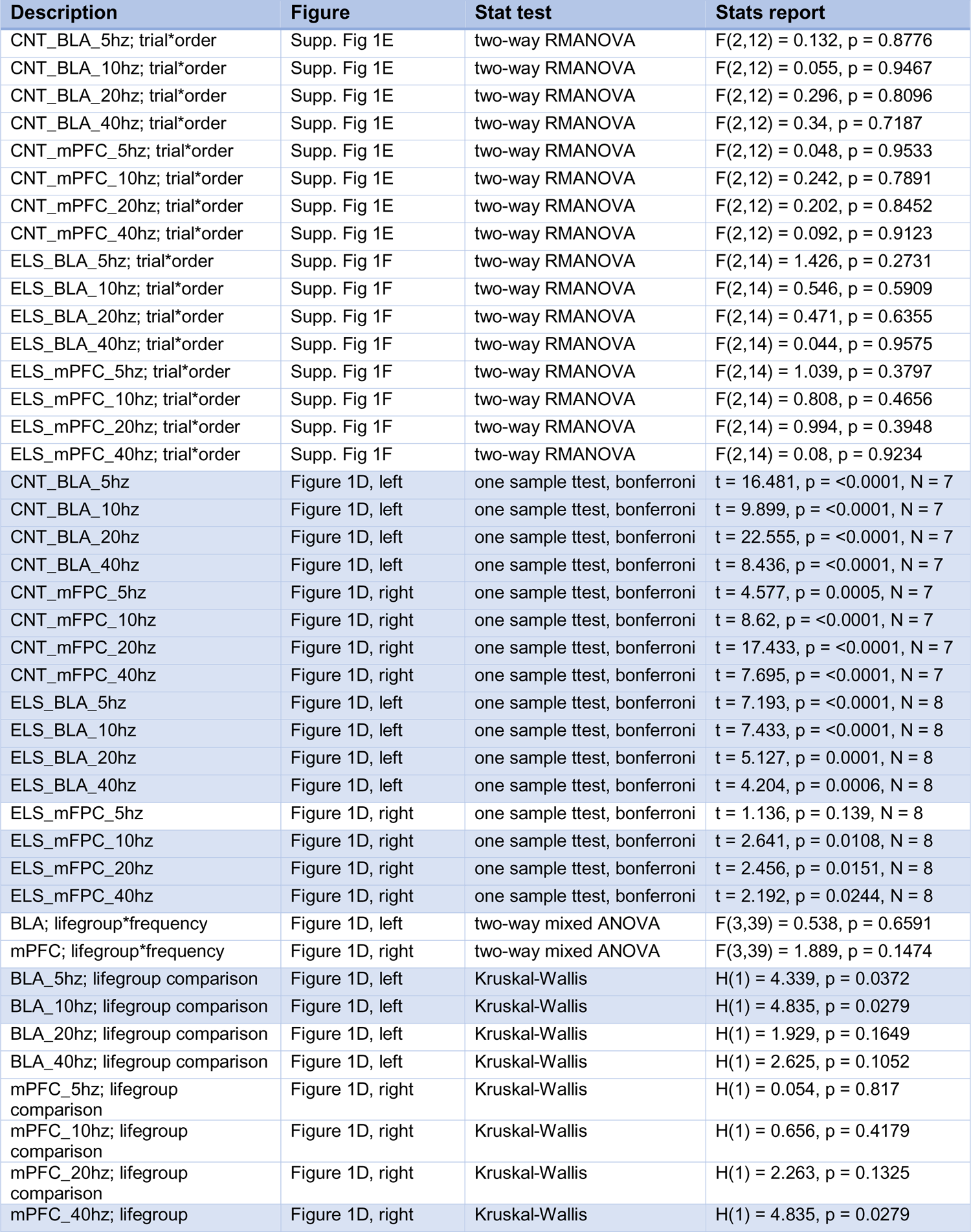
Highlighted rows: *p* < 0.05; Supp., Supplementary.

**Stats Table 2.** Highlighted rows: *p* < 0.05.

| Description | Figure | Stat test | Stats report |
| --- | --- | --- | --- |
| CNT_5hz | Figure 2E | one sample ttest, bonferroni | t = 1.9, p = 0.0364, N = 7 |
| CNT_10hz | Figure 2E | one sample ttest, bonferroni | t = 3.801, p = 0.0006, N = 7 |
| CNT_20hz | Figure 2E | one sample ttest, bonferroni | t = 4.306, p = 0.0002, N = 7 |
| CNT_40hz | Figure 2E | one sample ttest, bonferroni | t = 4.546, p = 0.0001, N = 7 |
| ELS_5hz | Figure 2E | one sample ttest, bonferroni | t = 0.831, p = 0.2189, N = 8 |
| ELS_10hz | Figure 2E | one sample ttest, bonferroni | t = 2.082, p = 0.0412, N = 8 |
| ELS_20hz | Figure 2E | one sample ttest, bonferroni | t = 4.818, p = 0.0016, N = 8 |
| ELS_40hz | Figure 2E | one sample ttest, bonferroni | t = 2.098, p = 0.0404, N = 8 |
| BLA; lifegroup*frequency | Figure 2E | two-way mixed ANOVA | F(3,75) = 0.769, p = 0.5152 |
| 5hz; lifegroup comparison | Figure 2E | Kruskal-Wallis | H(1) = 0.184, p = 0.6682 |
| 10hz; lifegroup comparison | Figure 2E | Kruskal-Wallis | H(1) = 0.735, p = 0.3914 |
| 20hz; lifegroup comparison | Figure 2E | Kruskal-Wallis | H(1) = 1.306, p = 0.2531 |
| 40hz; lifegroup comparison | Figure 2E | Kruskal-Wallis | H(1) = 4.592, p = 0.0321 |
| **Stats Table 2.** Highlighted rows: $p < 0.05$ | | | |

**Stats Table 3.** Highlighted rows: *p* < 0.05; Supp., Supplementary; N1, CNT; N2, ELS.

| Description | Figure | Stat test | Stats report |
| --- | --- | --- | --- |
| Fr. 20% input | Supp. Fig. 2A | Mann–Whitney U & BH correction | U=78.5, p=1.000, N1 = 13, N2 = 12 |
| Fr. 40% input | Supp. Fig. 2A | Mann–Whitney U & BH correction | U=91.0, p=0.991, N1 = 13, N2 = 12 |
| Fr. 60% input | Supp. Fig. 2A | Mann–Whitney U & BH correction | U=82.5, p=0.991, N1 = 13, N2 = 12 |
| Fr. Max input | Supp. Fig. 2A | Mann–Whitney U & BH correction | U=96.0, p=0.991, N1 = 13, N2 = 12 |
| Current Inj. at Half Max Fr | Supp. Fig. 2A | Mann–Whitney U & BH correction | U=82.5, p=0.991, N1 = 13, N2 = 12 |
| Input Resistance | Supp. Fig. 2A | Mann–Whitney U & BH correction | U=81.0, p=0.991, N1 = 13, N2 = 12 |
| IO slope | Supp. Fig. 2A | Mann–Whitney U & BH correction | U=65.0, p=0.991, N1 = 13, N2 = 12 |
| Max Fr. Rate | Supp. Fig. 2A | Mann–Whitney U & BH correction | U=89.0, p=0.991, N1 = 13, N2 = 12 |
| Resting Membrane Potential | Supp. Fig. 2A | Mann–Whitney U & BH correction | U=53.5, p=0.991, N1 = 13, N2 = 13 |
| Rheobase | Supp. Fig. 2A | Mann–Whitney U & BH correction | U=69.5, p=0.991, N1 = 13, N2 = 12 |
| AP amplitude | Supp. Fig. 2B | Mann–Whitney U & BH correction | U=69.0, p=0.773, N1 = 13, N2 = 12 |
| AP peak to trough | Supp. Fig. 2B | Mann–Whitney U & BH correction | U=66.0, p=0.773, N1 = 13, N2 = 12 |
| AHP amplitude | Supp. Fig. 2B | Mann–Whitney U & BH correction | U=101.0, p=0.773, N1 = 13, N2 = 12 |
| AP threshold | Supp. Fig. 2B | Mann–Whitney U & BH correction | U=87.0, p=0.773, N1 = 13, N2 = 12 |
| AP rise time | Supp. Fig. 2B | Mann–Whitney U & BH correction | U=74.0, p=0.849, N1 = 13, N2 = 12 |
| AP half width | Supp. Fig. 2B | Mann–Whitney U & BH correction | U=69.0, p=0.773, N1 = 13, N2 = 12 |
| Delta RMP (mV); lifegroup comparison | Figure 3D | independent t-test | t = 2.1573, p = 0.0437, N1 = 13, N2 = 9 |
| 80:20FR; lifegroup*treatment | Figure 3E | two-way mixed ANOVA | F(1,14) = 0.619, p = 0.4447 |
| Baseline; lifegroup comparison | Figure 3E | one-way ANOVA | F(1,14) = 0.294, p = 0.5962 |
| SCH; lifegroup comparison | Figure 3E | one-way ANOVA | F(1,14) = 5.868, p = 0.0296 |

**Stats Table 4.** Highlighted rows: *p* < 0.05; Supp., Supplementary.

| Description | Figure | Stat test | Stats report |
| --- | --- | --- | --- |
| CNT_SN; TH positive cell (avg count/slice) | Supp. Fig 3C | Pearson correlation | $r = 0.0189, p = 0.9195, N = 15$ |
| CNT_SN; Chrimson positive cell (avg count/slice) | Supp. Fig 3C | Pearson correlation | $r = -0.417, p = 0.0195, N = 13$ |
| ELS_SN; TH positive cell (avg count/slice) | Supp. Fig 3C | Pearson correlation | $r = -0.413, p = 0.0135, N = 19$ |
| ELS_SN; Chrimson positive cell (avg count/slice) | Supp. Fig 3C | Pearson correlation | $r = -0.087, p = 0.6191, N = 20$ |
| CNT_VTA; TH positive cell (avg count/slice) | Supp. Fig 3C | Pearson correlation | $r = -0.354, p = 0.0505, N = 15$ |
| CNT_VTA; Chrimson positive cell (avg count/slice) | Supp. Fig 3C | Pearson correlation | $r = -0.293, p = 0.1091, N = 13$ |
| ELS_VTA; TH positive cell (avg count/slice) | Supp. Fig 3C | Pearson correlation | $r = -0.593, p = 0.0001, N = 19$ |
| ELS_VTA; Chrimson positive cell (avg count/slice) | Supp. Fig 3C | Pearson correlation | $r = -0.349, p = 0.0395, N = 20$ |
| BLA nuclei; lifegroup comparison | Figure 4B | one-way ANOVA | $F(1,99) = 0.3636, p = 0.5479$ |
| BLA D1R1 density; lifegroup comparison | Figure 4C | one-way ANOVA | $F(1,99) = 2.212, p = 0.1401$ |
| Chrimson positive cells; lifegroup*region | Figure 4E, top | two-way ANOVA | $F(1,128) = 0.0141, p = 0.9057$ |
| SN-Chrimson positive cells; lifegroup comparison | Figure 4E, top | one-way ANOVA | $F(1,64) = 1.404, p = 0.2405$ |
| VTA-Chrimson positive cells; lifegroup comparison | Figure 4E, top | one-way ANOVA | $F(1,64) = 0.6578, p = 0.4203$ |
| TH positive cells; lifegroup*region | Figure 4E, bottom | two-way ANOVA | $F(1,128) = 0.0321, p = 0.8623$ |
| SN-TH positive cells; lifegroup comparison | Figure 4E, bottom | one-way ANOVA | $F(1,64) = 0.3167, p = 0.5755$ |
| VTA-TH positive cells; lifegroup comparison | Figure 4E, bottom | one-way ANOVA | $F(1,64) = 0.0194, p = 0.8896$ |
| SN-Chrim-TH cells; lifegroup comparison | Figure 4F | one-way ANOVA | $F(1,8) = 0.555, p = 0.1266$ |
| VTA-Chrim-TH cells; lifegroup comparison | Figure 4F | one-way ANOVA | $F(1,8) = 10.109, p = 0.0015$ |

**Stats Table 5.**
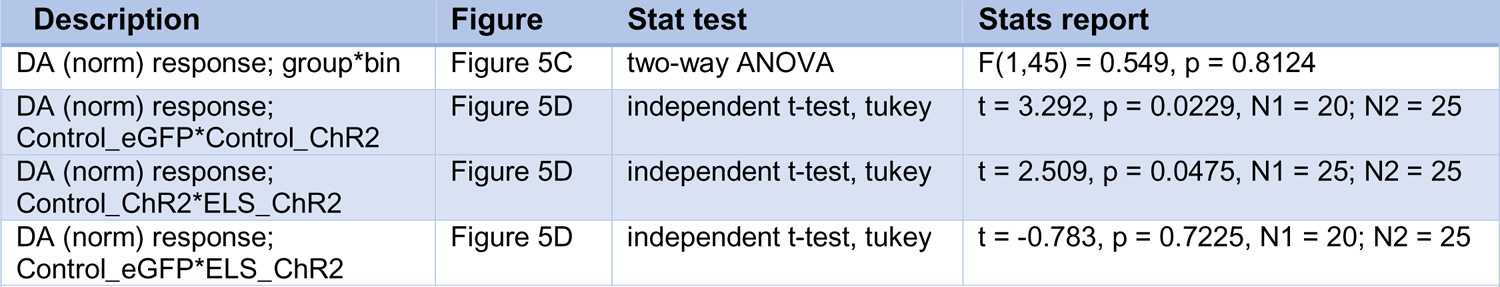
Highlighted rows: *p* < 0.05; N1, CNT; N2, ELS.

| Description | Figure | Stat test | Stats report |
| --- | --- | --- | --- |
| DA (norm) response; group*bin | Figure 5C | two-way ANOVA | $F(1,45) = 0.549$, $p = 0.8124$ |
| DA (norm) response;<br>Control_eGFP*Control_ChR2 | Figure 5D | independent t-test, tukey | $t = 3.292$, $p = 0.0229$, $N1 = 20$; $N2 = 25$ |
| DA (norm) response;<br>Control_ChR2*ELS_ChR2 | Figure 5D | independent t-test, tukey | $t = 2.509$, $p = 0.0475$, $N1 = 25$; $N2 = 25$ |
| DA (norm) response;<br>Control_eGFP*ELS_ChR2 | Figure 5D | independent t-test, tukey | $t = -0.783$, $p = 0.7225$, $N1 = 20$; $N2 = 25$ |
| **Stats Table 5.** Highlighted rows: $p < 0.05$; **N1**, CNT; **N2**, ELS | | | |

**Stats Table 6.** Highlighted rows: *p* < 0.05; N1, CNT; N2, ELS.

| Description | Figure | Stat test | Stats report |
| --- | --- | --- | --- |
| CNT_off_toy_toy | Figure 6E, left | one sample ttest, bonferroni | t = -0.615, p = 0.7264, N = 8 |
| CNT_on_toy_toy | Figure 6E, left | one sample ttest, bonferroni | t = -1.395, p = 0.9067, N = 8 |
| ELS_off_toy_toy | Figure 6E, left | one sample ttest, bonferroni | t = 0.69, p = 0.2508, N = 7 |
| ELS_on_toy_toy | Figure 6E, left | one sample ttest, bonferroni | t = -0.004, p = 0.5017, N = 7 |
| CNT_off_toy_female | Figure 6E, right | one sample ttest, bonferroni | t = 3.852, p = 0.0008, N = 8 |
| CNT_on_toy_female | Figure 6E, right | one sample ttest, bonferroni | t = 2.32, p = 0.0213, N = 8 |
| ELS_off_toy_female | Figure 6E, right | one sample ttest, bonferroni | t = 7.668, p = <0.0001, N = 7 |
| ELS_on_toy_female | Figure 6E, right | one sample ttest, bonferroni | t = 1.035, p = 0.1627, N = 7 |
| Toy-Toy; lifegroup*ChR2 | Figure 6E, left | two-way mixed ANOVA | F(1,13) = 0.787, p = 0.391 |
| Toy-Female; lifegroup*ChR2 | Figure 6E, right | two-way mixed ANOVA | F(1,14) = 5.155, p = 0.0395 |
| Toy-Toy; lifegroup comparison | Figure 6F | independent t-test | t = 1.187, p = 0.2583, N1 = 8; N2 = 7 |
| Toy-Female; lifegroup comparison | Figure 6F | independent t-test | t = 3.905, p = 0.0016, N1 = 8; N2 = 7 |
| **Stats Table 6.** Highlighted rows: $p < 0.05$; **N1**, CNT; **N2**, ELS | | | |

**Stats Table 7.**
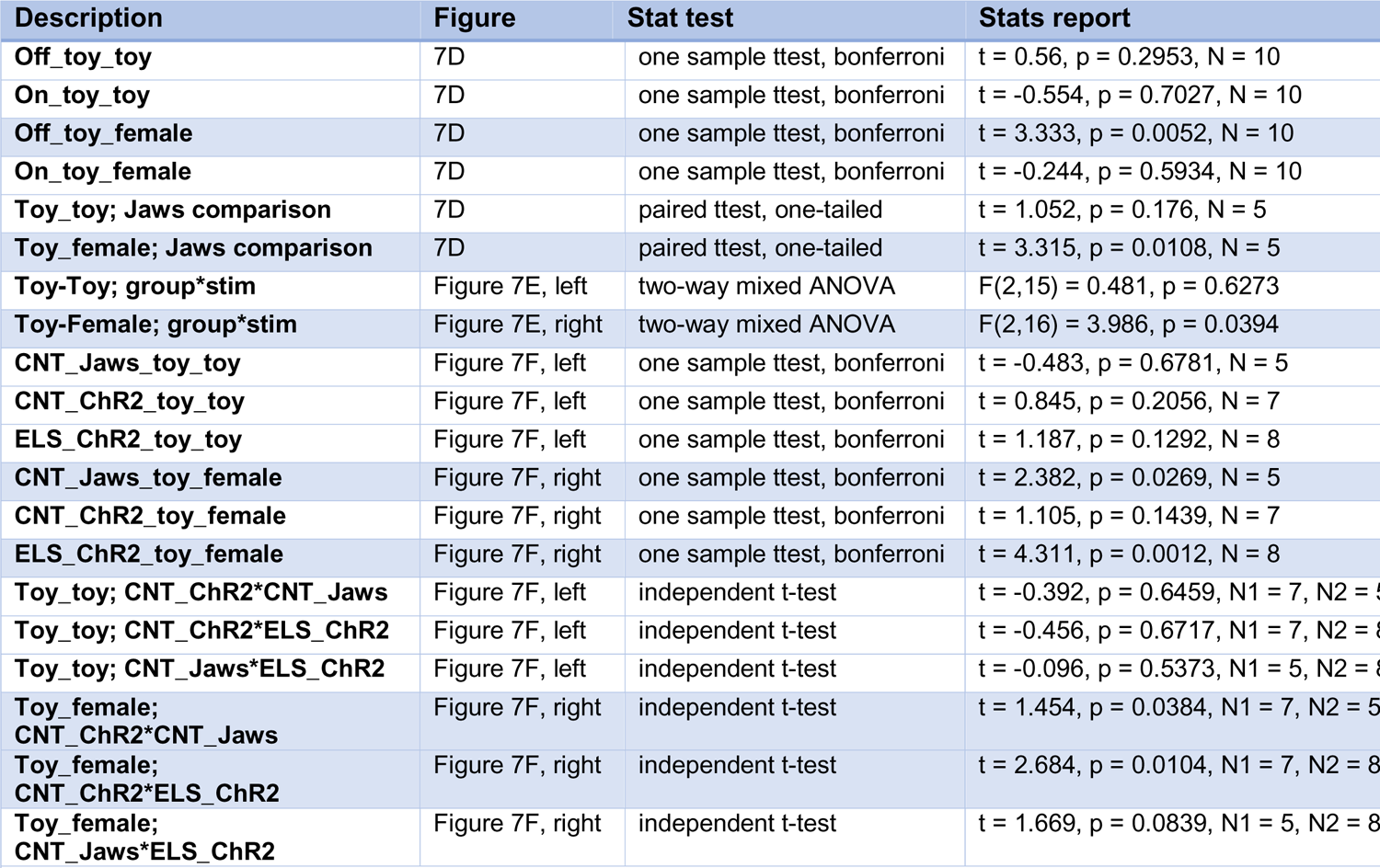
Highlighted rows: *p* < 0.05; N1, CNT; N2, ELS.

